# Management strategy evaluation for real-time closures in the short mackerel fishery in the Gulf of Thailand

**DOI:** 10.64898/2026.02.17.706503

**Authors:** Chonlada Meeanan, Pavarot Noranarttragoon, Piyachoke Sinanun, Wirat Sanitmajjaro, Takahashi Yuki, Methee Kaewnern, Matsuishi Takashi Fritz

## Abstract

Area and time restrictions are widely used in fisheries management for their simplicity and conservation benefits. Static closures (STCs) often fail to protect migratory fish; therefore, real-time closures (RTCs) are increasingly being adopted. However, RTCs require extensive and rapid data collection and analysis. The vessel monitoring system and daily landing reports provide near real-time data on fishing activities and fish abundance. We conducted a management strategy evaluation of RTCs and STCs in the Gulf of Thailand short mackerel fishery to clarify the efficacy of using RTCs to minimise fishing mortality, while considering their appropriate use with surveillance data for a migratory fish. The results support RTCs as more flexible and requiring a smaller closure area compared with STCs to achieve management objectives. We recommend gathering CPUE data on a monthly basis and using the highest CPUE threshold level to define a closure unit; no unit should be shut down until all units achieve the threshold level. Our results validate the efficacy of the RTC strategy for curtailing fishing mortality of a mobile species and demonstrate the effective use of RTCs to mitigate uncertainty in the migratory patterns of the species in an otherwise unpredictable, fluctuating environment.

## 1. Introduction

Time–area closures are an important fisheries management tool, widely used owing to their simple implementation and conservation benefits (Roberts and Polunin 1991; Arceo et al. 2013). This approach can reduce bycatch without significant loss of the target catch, especially for less migratory species (FAO 2003; Halpern et al. 2004); hence, area closures work best for species with limited movement patterns. Time–area closures are imposed in a specified area for a predetermined period (Munehara et al. 2021a). Static time–area closures (STCs) with fixed boundaries are simple to implement, yet may not accommodate changes in species distributions or fishing practices (Smith et al. 2021) and frequently fail to meet management goals (Bailey et al. 2010; Little et al. 2015; Munehara et al. 2021a), particularly for migratory species, which may form high-density areas (hotspots) in response to environmental, biological, and incidental factors (Dunn et al. 2011; Kongseng et al. 2020). Hence, imposing time–area closures to conserve a mobile species is generally challenging.

As a relatively recent innovation, dynamic or real-time closures (RTCs) are increasingly implemented in fisheries worldwide as an alternative to STCs (Needle and Catarino 2011). However, RTCs require extensive and rapid analysis of large volumes of data to inform decision-making (Bailey et al. 2010; Little et al. 2015). RTCs can be specifically targeted at zones exhibiting high fish abundance, areas where the catch predominantly comprises juvenile fish or sensitive species, or locations where a significant number of discards are expected based on catch composition (Kraak et al. 2014; Munehara et al. 2021a, 2021b).

Designing an RTC requires access to data and technological resources capable of providing real-time or predictive information. RTC management systems typically integrate information from multiple sources, such as onboard observers, at-sea inspections, monitoring systems, and enforcement tools like the vessel monitoring system (VMS) and daily landing reports. Holmes et al. (2011) report that effective identification of areas for closure often relies on the reliability and analysis of VMS data and the accuracy of logbook records. This information enables rapid responses to changing hotspot conditions. An example of an integrated approach is provided by Meeanan et al. (2023), who combined multiple sources of fisheries surveillance data—including VMS records, landing data, and fishing logbooks—to characterise the spatiotemporal distribution of fish, representing an initial effort to apply these data streams for near real-time monitoring of fishing fleet dynamics and fish abundance.

Dynamic fishery closure design is an integrative process that links data collection with the identification of fish density thresholds, operational rules, and management capacity. This includes defining a fish density threshold to trigger area closures, setting an appropriate closure period, ensuring timely and reliable data collection, applying effective analytical tools for decision-making, and implementing rules that are practical to enforce and compatible with existing fishing effort controls.

Establishing a fish density threshold, commonly expressed as catch per unit effort (CPUE), is a key step in defining areas of high fish abundance or hotspots for RTCs. The selected threshold plays an important role in achieving management objectives in that it determines when and where closures are triggered, and, consequently, the level of protection provided to the target species. In some fisheries, RTCs are implemented primarily to reduce bycatch or the capture of juvenile fish and are often regulated through catch quotas. Under such systems, hotspot thresholds can be defined simply as a specified proportion of the catch quotas being reached or exceeded (Holmes et al. 2011; Little et al. 2015). Therefore, identifying an appropriate hotspot threshold is a critical component of effective real-time spatial management because it directly impacts decision-making, resource allocation, and response effectiveness (Little et al. 2015).

The existing literature has not adequately addressed how to gauge the information-gathering period used to determine hotspot locations for RTCs, even though the effectiveness of management hinges on clearly defined ‘closure rules’. RTCs depend on rapid data to track the movement of target fish within fishing zones, to adjust fishing regulations, and safeguard marine species at the most favourable time and place, especially considering unpredictable shifts in distribution. Consequently, when determining hotspots of fish abundance, it is imperative to consider the data-collection period relative to the typical scale of fish movement within the studied ecosystem. For instance, in the Scottish North Sea demersal fishery, where prior research minimised cod mortality by using 14 days of historical fishing vessel position data from VMS and logbooks to evaluate the CPUE hotspot, the “triggered area” was thereby defined using CPUE from the same calendar month in the previous year (Holmes et al. 2011; Little et al. 2015).

Implementing RTCs in effort-controlled fisheries is inherently complex. When fishing effort is regulated through measures such as days-at-sea, defining an appropriate CPUE threshold to identify high-density hotspots for area closures is particularly challenging. The choice of this threshold directly affects the size and duration of a closure and, consequently, the fishing mortality of target species. Because RTCs rely heavily on dynamic and responsive decision rules, their design and implementation require careful evaluation in advance to understand their potential impacts, limitations, and feasibility in achieving management objectives.

Management strategy evaluation (MSE) provides a formal and systematic framework for testing fisheries management measures under uncertainty. MSE involves using simulation of the entire management cycle—linking stock dynamics, monitoring, assessment, and decision-making—to allow different strategies to be compared in a repeatable way. Through this approach, managers can evaluate trade-offs and robustness before real-world implementation. In this study, MSE was applied to test alternative time–area closure strategies, namely STCs and RTCs, with the aim of assessing whether dynamic closures can more effectively reduce fishing mortality than static measures, and whether RTCs can be feasibly implemented using surveillance-derived information.

The application of RTCs was examined in the context of the fishery for short mackerel *Rastrelliger brachysoma*, a highly migratory and commercially important pelagic species distributed in the Gulf of Thailand and throughout Southeast Asia. Short mackerel migrate across national boundaries, making the species a suitable case for evaluating dynamic spatial management strategies. Since 2018, STCs have been implemented to protect this species in the Gulf of Thailand (Fig. 1); however, the effects on fishing mortality and the sensitivity of STCs to this highly mobile species remain unclear.

**Fig. 1.**
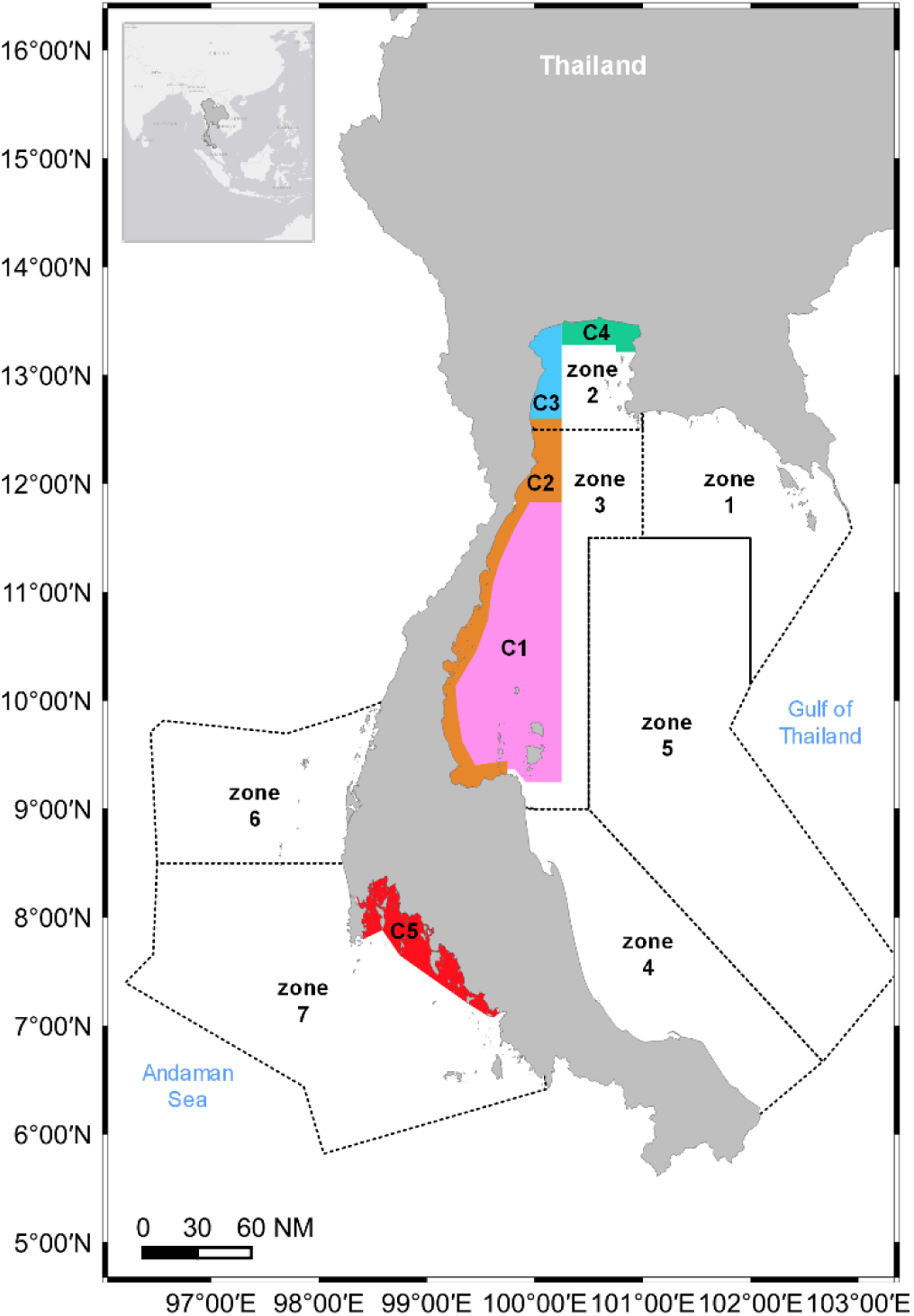
The Gulf of Thailand (GOT) and the Andaman Sea (ADS) are two distinct fishing grounds governed as separate management areas in Thailand. The entire fisheries statistical zone is divided into seven zones: Eastern GOT (zone 1), Upper GOT (zone 2), Western GOT (zone 3), Southern GOT (zone 4), Middle GOT (zone 5), Upper ADS (zone 6), and Lower ADS (zone 7).

This study applied a MSE framework to the short mackerel purse-seine fishery in the Gulf of Thailand with the objective of evaluating the potential of RTCs as a management tool for a transboundary, migratory species. Specifically, we aimed to assess and compare the effectiveness of STCs versus RTCs in reducing fishing mortality, and to identify an optimal RTC decision rule for the short mackerel fishery in the Gulf of Thailand. To achieve this, our three main objectives were to: (1) investigate the temporal coverage of data applied in identifying CPUE hotspots; (2) determine the CPUE threshold used to designate closure units; and (3) compare the performance of STCs versus RTCs in the short mackerel purse-seine fishery in the Gulf of Thailand in 2019–2020.

## 2. Materials and methods

### 2.1. Simulation approach

The MSE framework applied in this study consisted of two main components, as illustrated in Fig. 2: operating models that represent realistic fishery and stock processes, and a management process that applies alternative time–area closure strategies to these operating models. Together, these components simulate how management decisions interact with underlying population dynamics and allow the performance of different management scenarios to be evaluated in a controlled and repeatable manner.

**Fig. 2.**
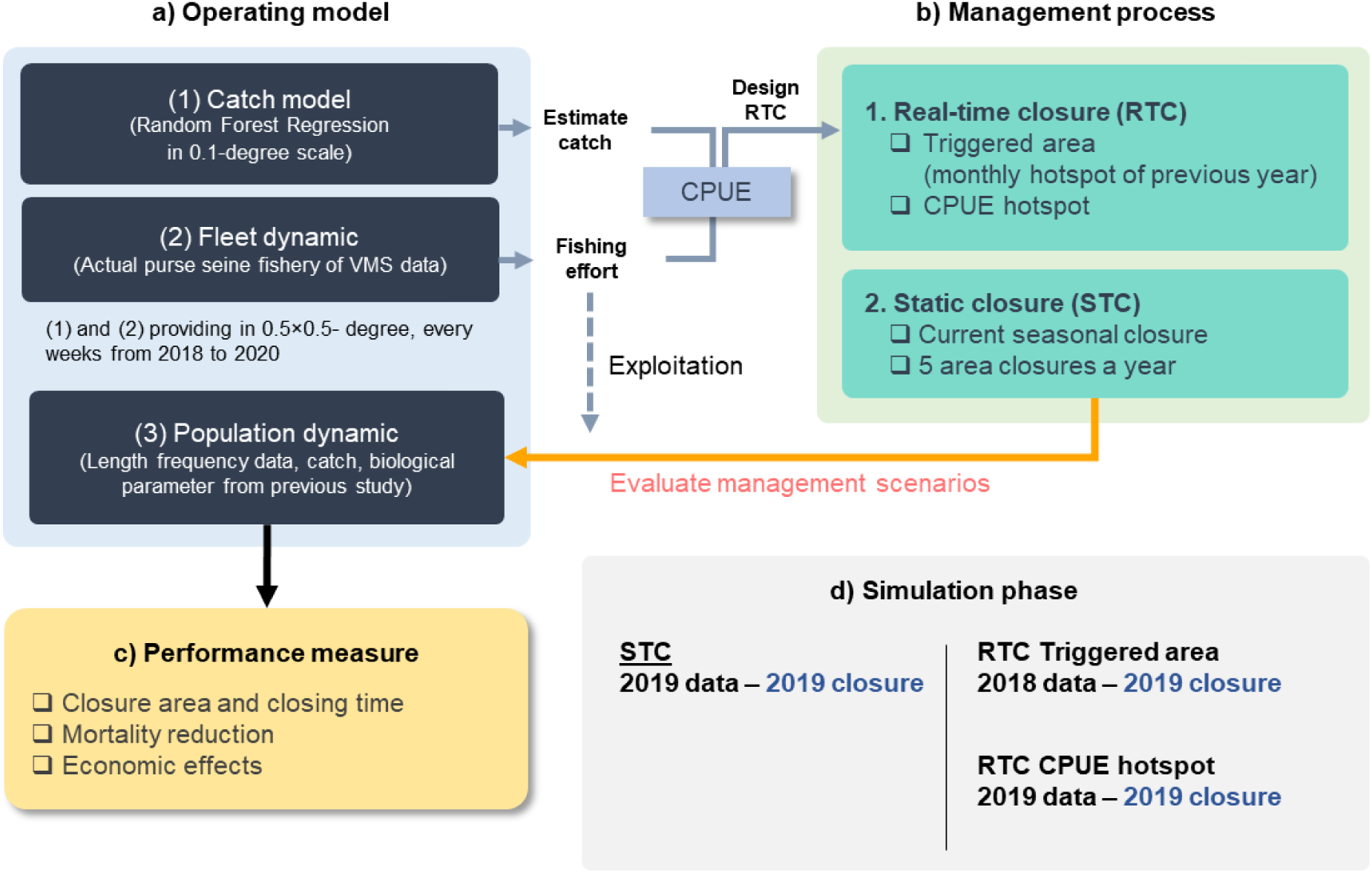
Diagram of the management strategy evaluation (MSE) framework. a) The operating model comprised a catch model representing realistic spatial catch distributions, and a fleet dynamics model based on observed purse-seine vessel monitoring system (VMS) data, together with short mackerel population dynamics informed by length frequency, catches, and biological parameters. b) The evaluation considered two alternative management strategies: real-time closure (RTC) versus static closure (STC). c) Management performance was assessed across three domains: closure area and duration; reduction in fishing mortality; and economic effects. d) The simulation phase depicts the temporal alignment and duration of the datasets used to trigger and evaluate fishery closures within the MSE simulations.

The operating models (Fig. 2a) consisted of two interacting components—the catch component and fleet dynamics component. The catch model estimates spatial patterns of catch, while the fleet dynamics model represents fishing activity using information derived from VMS data. Estimated catch and fishing effort were combined on a weekly basis to derive CPUE, which was used as an indicator of fish abundance and served as the primary input for triggering an RTC in the simulations. The management process (Fig. 2b) evaluates two alternative strategies: RTC and STC. These strategies were applied to the operating models, and the population dynamics of short mackerel in the Gulf of Thailand were then simulated to examine how the stock would respond over time under different closure designs.

Management performance was evaluated across three domains (Fig. 2c): the spatial and temporal characteristics of closures, reductions in fishing mortality, and economic effects. Economic effects were assessed indirectly through changes in population dynamics, using information on length frequency distributions, catches, and biological parameters from previous studies. The simulation phase (Fig. 2d) defined the temporal structure of data use for each scenario, enabling assessment of how different data periods and triggering rules might influence management outcomes. For STC, data from 2019 were used to implement and evaluate closures within the same year. For the RTC-triggered-area scenario, data from 2018 were used to define closure triggers, with the subsequent closures then applied in 2019. For the RTC-CPUE hotspot scenario, 2019 data were used both to identify hotspots and to implement closures during 2019.

### 2.2. Operating model

#### 2.2.1. Catch model

Catch distribution during the simulation period (2018–2019) was characterised using CPUE as an indicator of the probability of fish occurrence, derived from integrated fisheries surveillance data, including VMS records, landing data, and fishing logbooks. The methodology for estimating the spatiotemporal CPUE distribution followed Meeanan et al. (2023), who introduced the use of surveillance data for near real-time assessment of short mackerel abundance. This established approach was applied to the 2018–2019 dataset to describe spatial and temporal patterns of short mackerel distribution relevant to the present analysis.

Because the data used to characterise fish distribution were collected during a period when seasonal closures were in place, fish occurrence within the closed areas could not be directly observed. To overcome this limitation and to estimate fish distribution in the areas not accessible to fishing under existing time–area restrictions, an additional modelling approach, random forest regression (RFR), was applied. Catch data from the Gulf of Thailand during 2018–2019 were used as a training dataset, while environmental variables were included as predictors, following Meeanan et al. (2023), who introduced the use of spatial and environmental covariates for assessments of short mackerel abundance. This modelling framework enabled the prediction of weekly catch distributions in data-limited areas, providing a more complete spatial representation of short mackerel distribution for the subsequent MSE.

The potential distribution of catch in areas with missing observations was estimated on a weekly basis, using a predictive model in which catch was expressed as a function of environmental and spatial variables:

Catch = SST + Chl 𝑎 + Sea depth + Distance from shore + Latitude + Longitude where Catch was defined as the estimated catch (kg) within each 0.03° × 0.03° spatial grid. Sea surface temperature (SST, °C) was derived from 8-day composite products, and chlorophyll-*a* concentration (mg m⁻³) was obtained at 4-km spatial resolution from MODIS-Aqua/Terra data. Sea depth (m) was sourced from the British Oceanographic Data Centre at a spatial resolution of 15 arc-seconds (∼0.004°). Distance from shore for each VMS position was calculated as the shortest distance to the coastline, using established serial programming routines (Stack Overflow 2014). The vessel locations, including latitude and longitude, were collected from the VMS on an hourly basis. The weekly estimated catch was gathered in the management unit (Fig. 3) and used for the preparation of possible RTC scenarios.

**Fig. 3.**
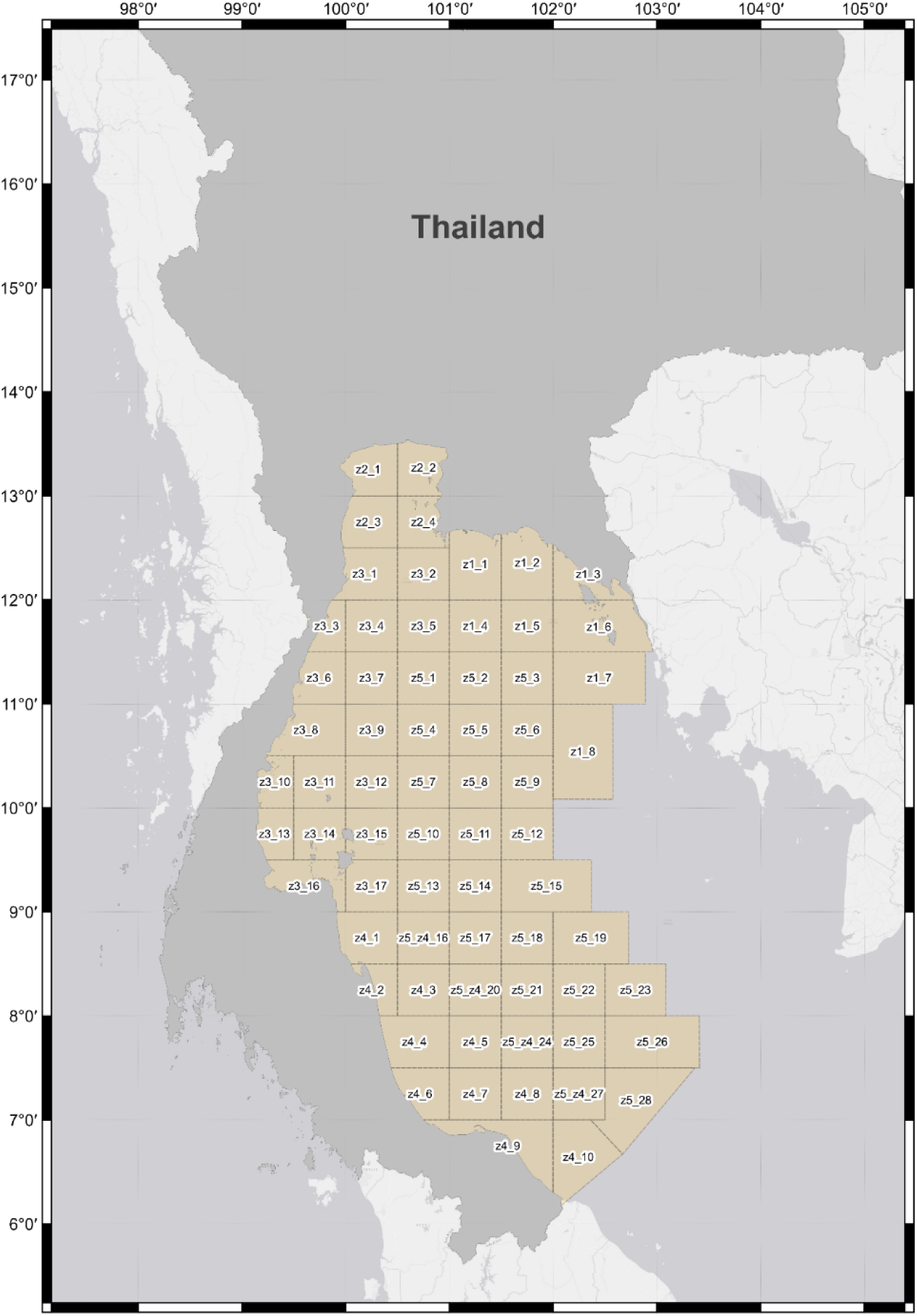
The management units designated for real-time closures in the short mackerel fishery are specific locations in the Gulf of Thailand that have been partitioned into roughly 0.5 × 0.5-degree grid units; for areas adjacent to the land or the ocean boundary, the ratio of one side remains constant at 0.5 degrees, notwithstanding potential variations in the unit sizes. The letter ‘z’ for zone is followed by a number identifying the designated zone under the current statistical management zone framework used in Thailand, which encompasses a total of five distinct zones in the gulf; the number after the underscore denotes the sequential arrangement of units established in the context of analysis.

#### 2.2.2. Fishing effort

The weekly fishing effort values were obtained based on the actual data (Meeanan et al. 2023) and used in the operating model. Some locations and times are unavailable because of the present time–area closure setting. The RFR prediction model was used to estimate fishing effort, as follows:

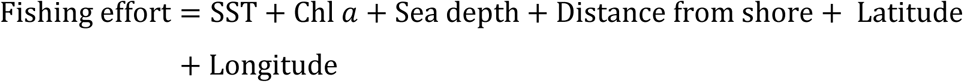

where fishing effort is defined as the number of fishing operations that were undertaken in a 0.03° × 0.03° square during a given number of hours, and the other variables were described from the same procedure as the catch prediction variables. Finally, fishing effort was predicted at a 7-day temporal resolution, with outputs at a 0.03° × 0.03° spatial resolution subsequently standardised to the management units (Fig. 3) and then used for the preparation of possible RTC scenarios.

Next, the CPUE of a specific management unit 𝑚 at a specific time *t* was calculated as:

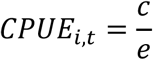

where 𝑐 is the sum of the estimated catch (kg) in management unit 𝑖, and 𝑒 is the amount of fishing activity (days) in management unit 𝑖.

### 2.3. Management scenarios: STC versus RTC

One STC scenario and 18 RTCs scenarios were evaluated (Table 1), and their performances compared.

**Table 1.** The time–area closure management scenarios (STC = static closure; RTC = dynamic, real-time closure) used for management strategy evaluation of the short mackerel fishery in the Gulf of Thailand (GOT). 3SD and 4SD denote 3 or 4 standard deviations from the mean catch per unit effort (CPUE), respectively.

| Scenario no. | Management scenario | Closure scenario | Data collection range | CPUE threshold level | Mortality reduction |
| --- | --- | --- | --- | --- | --- |
| 1 | STC | Static closure following the existing five seasonal closures implemented in the GOT (see Fig. 1) |  |  |  |
| 2 | RTC | RTC_my_3SD | Triggered area | 3SD | Close for 1 month |
| 3 | RTC | RTC_my_4SD | Triggered area | 4SD | Close for 1 month |
| 4 | RTC | RTC_mm_3SD_5% | Monthly CPUE | 3SD | 5% |
| 5 | RTC | RTC_mm_3SD_10% | Monthly CPUE | 3SD | 10% |
| 6 | RTC | RTC_mm_3SD_20% | Monthly CPUE | 3SD | 20% |
| 7 | RTC | RTC_mm_3SD_30% | Monthly CPUE | 3SD | 30% |
| 8 | RTC | RTC_mm_4SD_5% | Monthly CPUE | 4SD | 5% |
| 9 | RTC | RTC_mm_4SD_10% | Monthly CPUE | 4SD | 10% |
| 10 | RTC | RTC_mm_4SD_20% | Monthly CPUE | 4SD | 20% |
| 11 | RTC | RTC_mm_4SD_30% | Monthly CPUE | 4SD | 30% |
| 12 | RTC | RTC_2w_3SD_5% | Biweekly CPUE | 3SD | 5% |
| 13 | RTC | RTC_2w_3SD_10% | Biweekly CPUE | 3SD | 10% |
| 14 | RTC | RTC_2w_3SD_20% | Biweekly CPUE | 3SD | 20% |
| 15 | RTC | RTC_2w_3SD_30% | Biweekly CPUE | 3SD | 30% |
| 16 | RTC | RTC_2w_4SD_5% | Biweekly CPUE | 4SD | 5% |
| 17 | RTC | RTC_2w_4SD_10% | Biweekly CPUE | 4SD | 10% |
| 18 | RTC | RTC_2w_4SD_20% | Biweekly CPUE | 4SD | 20% |
| 19 | RTC | RTC_2w_4SD_30% | Biweekly CPUE | 4SD | 30% |

#### 2.3.1. STC scenario

Five seasonal STCs are imposed in the seven statistical management zones in the Gulf of Thailand (GOT) and the Andaman Sea (ADS), depicted in Fig. 1 in terms of their geographical extent and lengths of time. The first closure (C1) is implemented annually in the central GOT from 15 February to 15 May. From 16 May to 14 June, the second closure (C2) reduces the closed area to within ∼7 nautical miles from the shore and the northern portion of the initial closure zone. The third (C3) and fourth (C4) closures are established in the inner GOT from 15 June to 15 August on the western side and from 1 August to 30 September on the northern side, respectively. In the ADS, a single closure (C5) is enforced from 1 April to 30 June.

#### 2.3.2. RTC scenarios

To assess the sensitivity of management outcomes to different RTC design choices, a suite of RTC scenarios was evaluated that varied in (1) the temporal range for identifying CPUE hotspots, (2) the CPUE threshold applied to define closure units, and (3) the targeted level of fishing mortality reduction (Table 1, Fig. 4). These scenarios were designed to examine how alternative RTC configurations would influence management performance, as described below.

(1) Temporal range of data collection for identifying CPUE hotspots:

**Fig. 4.**
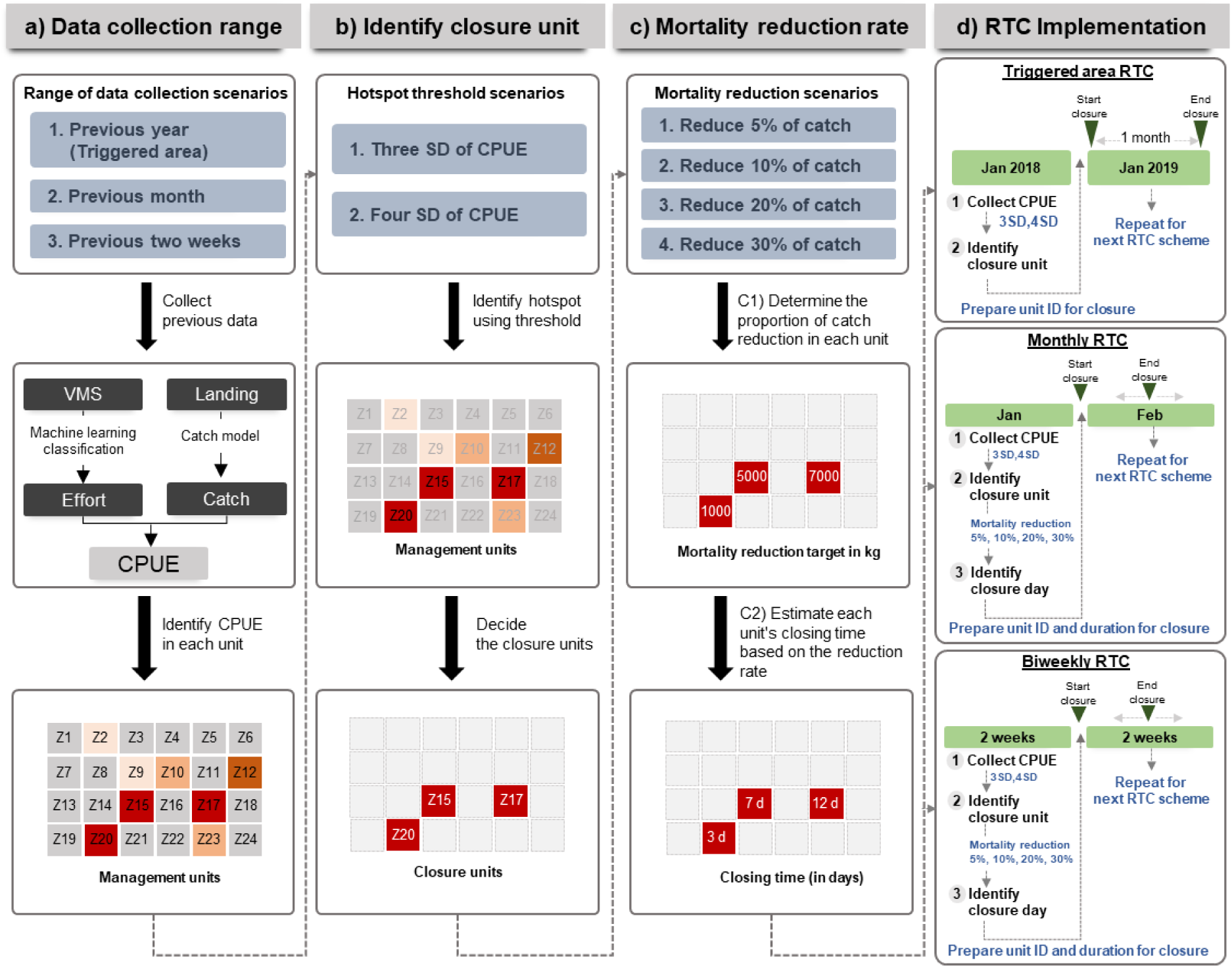
Diagram of RTC implementation for the management scenarios listed in Table 1, including: a) assumptions about the data-collection interval for identifying CPUE and hotspots; b) the potential of CPUE data for defining a closure area using different CPUE threshold levels; and c) target levels of mortality reduction. d) The process of implementing an RTC after defining the unit and closure time. The letters ‘z’, ‘ID’, and ‘SD’ denote zone, unit identification, and standard deviation, respectively.

**Fig. 5.**
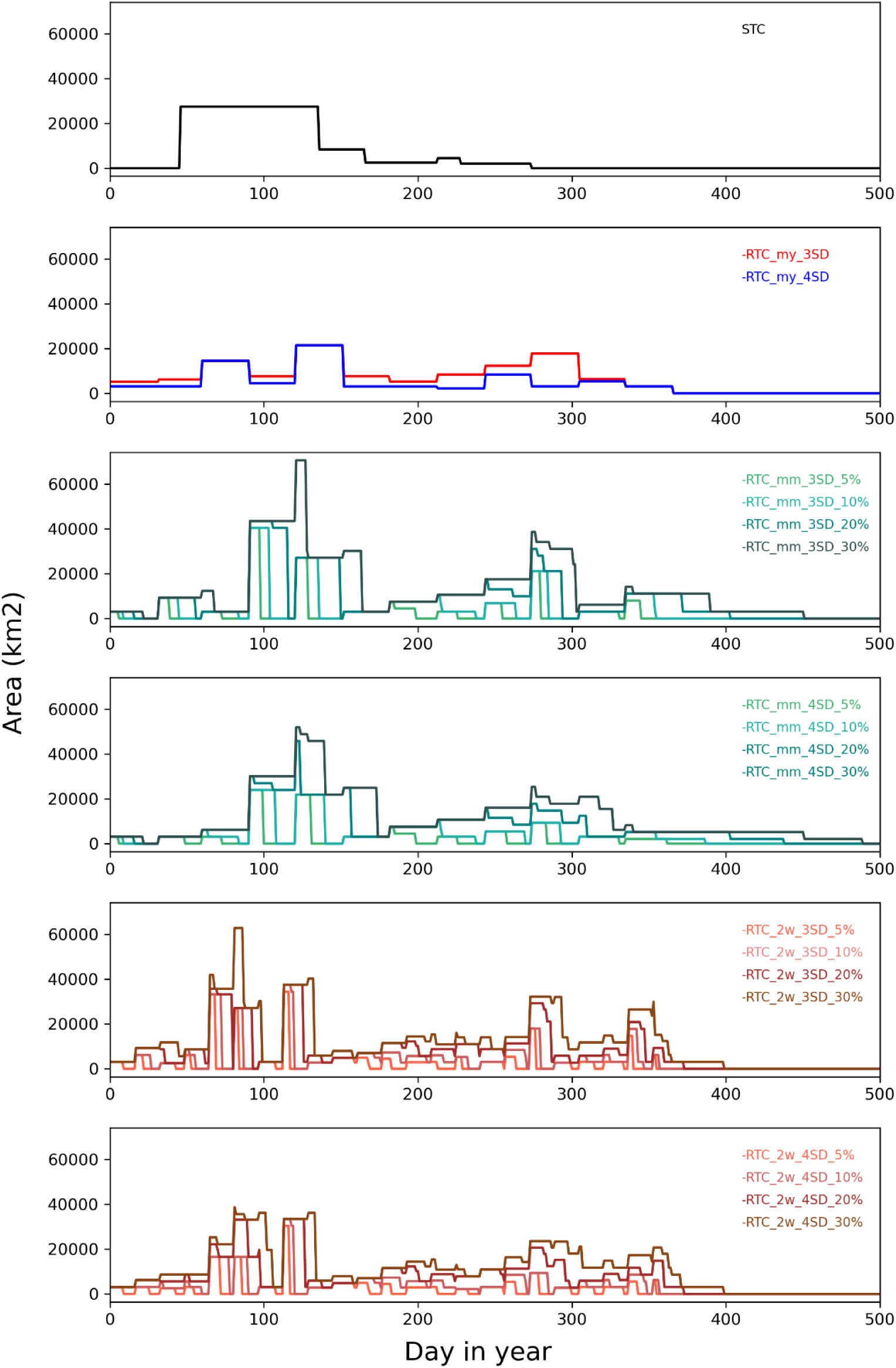
Temporal distribution of the fishery closure areas (km^2^) in the Gulf of Thailand, with daily resolution over the 2019–2020 period (Day 1 of 2019 to Day 500), under each management scenario. STC = static closure; RTC = dynamic, real-time closure.

Three RTC scenarios were evaluated to examine how the length of the surveillance data used to identify CPUE hotspots influences closure performance (Table 1, Fig. 4). The scenarios differed only in the time window of data used to define hotspots, thereby reflecting uncertainty in the spatial and temporal movements of fish.

(i) Triggered area: Hotspots were identified using surveillance data from the same month in the previous year.
(ii) Monthly CPUE closure area: Hotspots were determined using data collected 1 month prior to the closure decision.
(iii) Biweekly CPUE closure area: Hotspots were identified using data collected 2 weeks prior to the closure decision.

(2) The CPUE threshold level for defining a closure area:

Two CPUE threshold scenarios were tested to examine how the definition of a hotspot influences a closure decision (Table 1, Fig. 4). The scenarios differed only in the stringency of the CPUE threshold used to trigger a closure, as follows:

(i) Three standard deviations of CPUE (3SD): A management unit was classified as a hotspot when its CPUE exceeded 3 standard deviations (SD) above the annual mean CPUE.
(ii) Four standard deviations of CPUE (4SD): A more restrictive threshold of 4 standard deviations (SD) was applied.

In both scenarios, management units meeting or exceeding the threshold were selected for closure; if no unit exceeded the threshold during a given period, the unit with the highest CPUE was designated for closure.

(3) Targeted level of mortality reduction:

RTC scenarios also differed in the target level of fishing mortality reduction (Table 1, Fig. 4c). Four mortality reduction levels were evaluated: 5%, 10%, 20%, and 30%, representing increasing management stringency. The percentage of mortality reduction was calculated using the proportion of historical catch data gathered before the closure decision.

For example, catch and CPUE data from January were used to determine closures for February. If the total catch in January was 60,000 kg and the target mortality reduction was set at 5%, the total reduction target for the subsequent period was 3,000 kg. This reduction target was then allocated among the management units selected for closure in February in proportion to their CPUE levels. Because multiple management units may be closed within a given period and each unit has a different CPUE, the mortality reduction assigned to each unit was weight by its relative contribution to total CPUE (Fig. 4c), defined as:

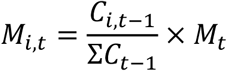

where 𝑀_𝑖,𝑡_ is the mortality reduction in kg allocated to management unit 𝑖 during closure period 𝑡; 𝐶_𝑖,𝑡−1_ is the CPUE of unit 𝑖 in the preceding period; Σ𝐶_𝑡−1_ is the sum of CPUE across all management units selected for closure; and 𝑀_𝑡_ is the total target mortality reduction for the period.

To ensure that RTCs achieve the predefined mortality reduction target, the closure duration for each management unit was determined based on its allocated mortality reduction and CPUE (Fig. 4c). Closure duration was calculated as:

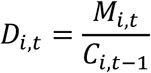

where 𝐷_𝑖,𝑡_ represents the number of closure days for management unit 𝑖. This approach ensures that the combined effect of closures across all units within a given period corresponds to the intended overall reduction in fishing mortality.

Implementation of the different RTC schemes were carried out after determining the CPUE in each management unit, deciding the closure units, and calculating the closure times based on the targeted level of mortality reduction in each unit (Fig. 4d). As an exception, the mortality reduction level was not defined for the triggered RTC scenario. Consequently, all units were scheduled to be closed for the duration of 1 month, beginning in the following month.

### 2.4. Performance evaluation

The effectiveness of closure scenarios was determined by the total closure area, the actual mortality reduction, and the economic effect resulting from closure (Table 2). To further clarify the application of different RTC schemes, the appropriateness of the data-gathering range and CPUE threshold were addressed.

**Table 2.**
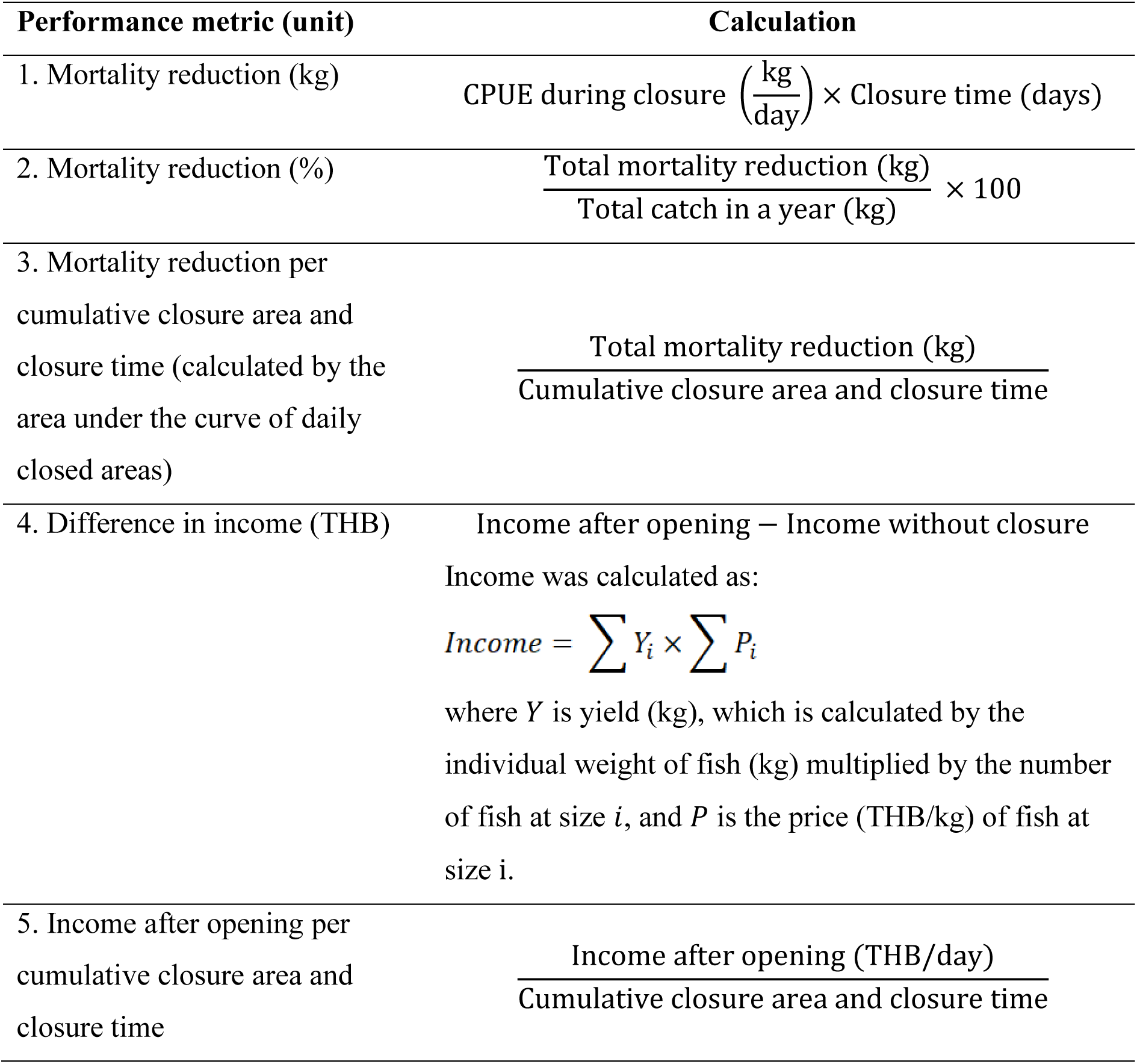
Summary of the performance of closure scenarios for the short mackerel fishery in the Gulf of Thailand. THB = Thai Baht.

A graph was generated to examine the everyday proportion of closed areas by plotting the closed area against every single day within a year. To analyse the cumulative closure area and closure time, the area under the graph was used to describe aggregating closure areas against 1 day in 2019. This was carried out by applying the trapezoidal rule to the average values of the replicates at each time point (Scheff et al. 2011). Next, the cumulative closure area and closure time were calculated to compare temporal changes in the closure areas.

The population dynamic model was used to analyse the economic effects before the closure schemes. The exploitation of fish stocks was assessed through the exponential decay model (Sparre and Venema 1998) together with the von Bertalanffy growth equation, to investigate changes in fish mortality and survival across time under different closure scenarios:

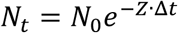

where 𝑁_𝑡_ is the number of surviving target fish after time 𝑡 (1/year), 𝑁_0_ is the initial number of fish, and 𝑍 is the total instantaneous mortality rate in units of time (1/year). 𝑍 was calculated as 𝑍 = Fishing mortality (𝐹) + Natural mortality (𝑀), and *Δi* is the time difference (1/year) between 𝑁_0_ and 𝑁_𝑡_.

Thailand’s Department of Fisheries provided length frequency data for short mackerel gathered in 2019. These data were divided monthly into statical management zones (Fig. 1) to calculate the number of fish survivors over time using the exponential decay model. For assessing this model, the length data were converted to weight and age using the length–weight relationship equation and the inverse von Bertalanffy growth equation (Sparre and Venema 1998), using the biological parameters presented in Table 3.

**Table 3.** Biological parameters of short mackerel *Rastrelliger brachysoma* in the Gulf of Thailand.

| Parameters | Value | Reference |
| --- | --- | --- |
| Intercept ($a$) | 0.0120 | Sritakon et al. (2011) |
| Slope ($b$) | 2.9968 | Sritakon et al. (2011) |
| $L_{\infty}$ (cm) | 23.36 | Prasertsook (2021) |
| $K$ (1/year) | 1.6 | Prasertsook (2021) |
| Natural mortality ($M$) (1/year) | 1.79 | Prasertsook (2021) |
| $t_0$ (1/year) | -0.2346 | Prasertsook (2021) |

The length–weight relationship equation is expressed as:

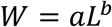

where 𝑊 is the body weight (g), 𝐿 is the standard body length (cm), 𝑎 is the intercept, and 𝑏 is the slope.

The inverse von Bertalanffy growth equation was calculated as:

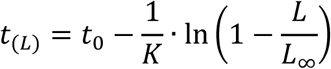

where 𝑡_(𝐿)_ is age at length 𝐿; 𝑡_0_ is the theoretical age at which length is zero; 𝐾 is the growth rate coefficient; and 𝐿_∞_ is the asymptotic average length.

The appropriateness of the data-gathering range for determining the CPUE under each scenario was investigated using different values of CPUE, as follows:

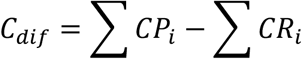

where 𝐶_𝑑𝑖𝑓_ is the difference in CPUE; 𝐶𝑃 is the predetermined CPUE applied to the RTC in unit 𝑖; and 𝐶𝑅 is CPUE during the closure of unit 𝑖.

## 3. Results

### 3.1. Closure area and duration

Fig. 4 depicts variations in the total closure areas and closure times under the different management scenarios. In general, the STC scenario entailed consistent closure of the short mackerel fishery within specific time-frames, whereas the RTC scenario entailed closures that varied spatially and temporally throughout the year. Fig. 6 shows a comparison of the management scenarios in terms of the estimated cumulative closure areas and closure times, as obtained by summing the closure areas over a 1-day interval. The cumulative closure area and closure time of an RTC was comparatively lower than that of the STC in some scenarios, particularly under mortality reduction targets of 5% or 10%. Under RTC scenarios exhibiting greater cumulative closure areas and closure times, the mortality reduction targets increased to 20% and 30%.

**Fig. 6.**
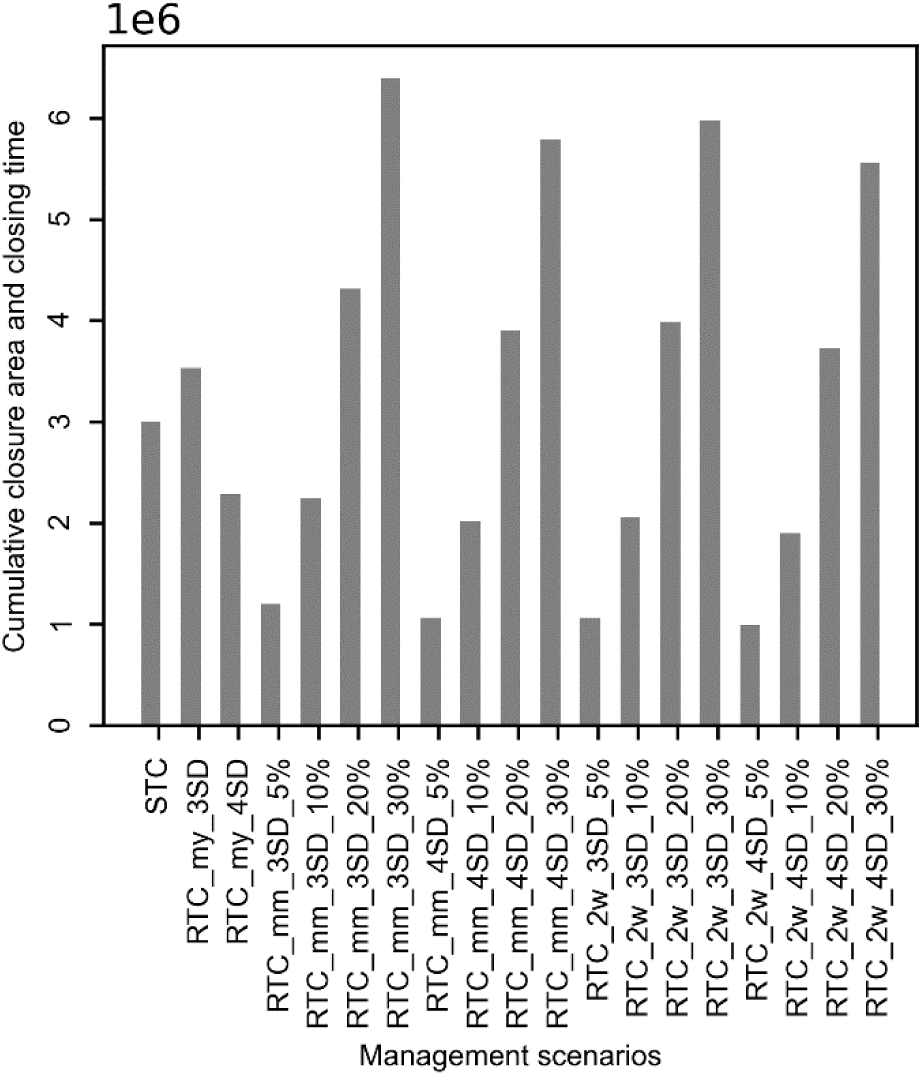
Comparison of cumulative closure area and closure time, estimated by calculating the area under the graph by aggregating the closed areas against a 1-day interval in each 2019 and 2020, under various management scenarios. STC = static closure; RTC = dynamic, real-time closure.

### 3.2. Mortality reduction

The RTC strategy showed greater potential to minimise fishing mortality of short mackerel than the STC strategy. The RTC scenarios displayed a clear capacity to reduce fishing mortality especially as the mortality reduction target percentage was increased (Fig. 7). Under RTC scenarios designed to reduce fishing mortality by 5%, 10%, 20%, or 30%, the actual percentage mortality after opening was below the intended level. The RTCs achieved consistently higher reductions in fishing mortality per cumulative closure area and closure time (Fig. 7) than the STC, across all scenarios.

**Fig. 7.**
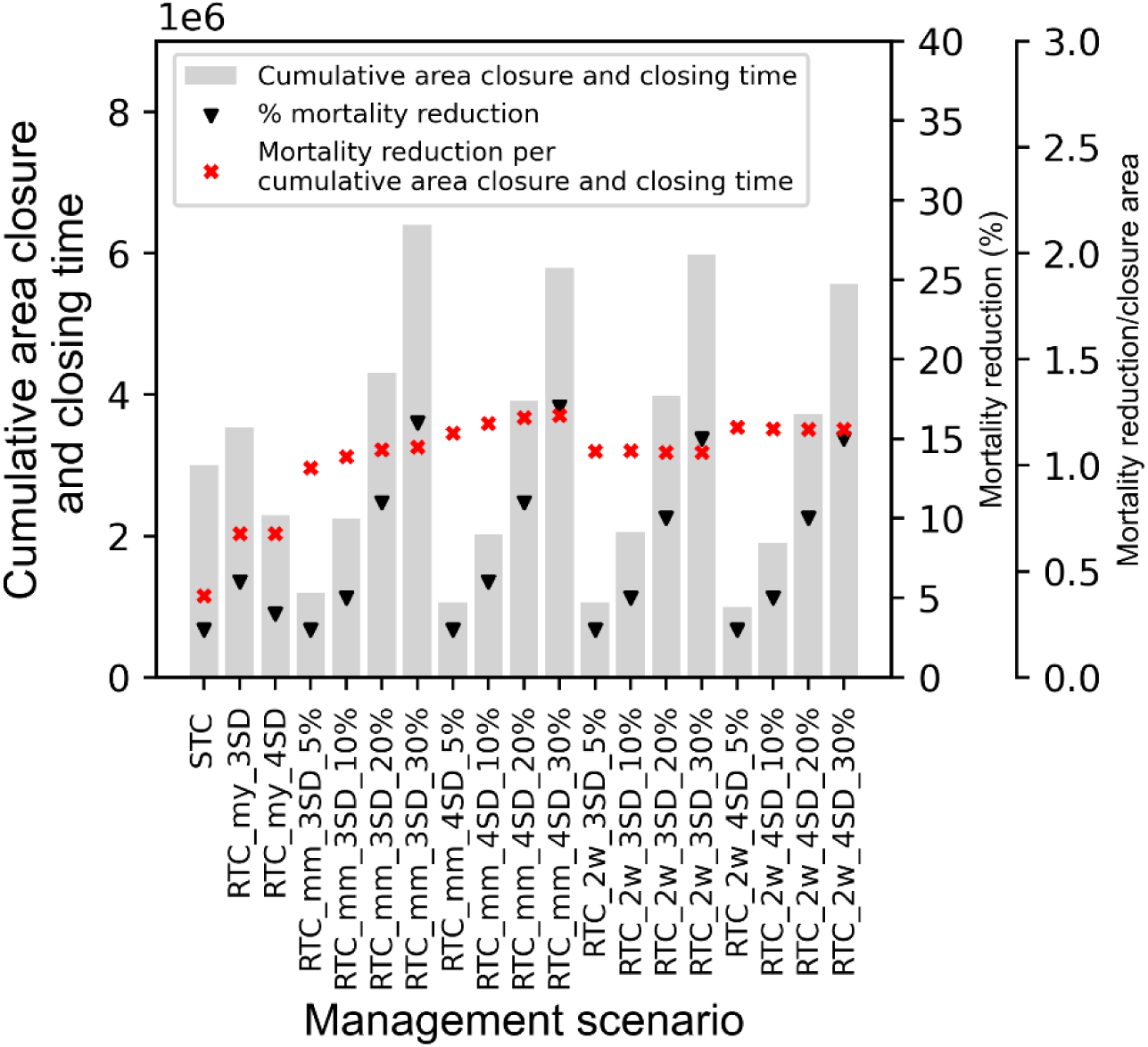
Performance metrics referring to the reduction of fishing mortality under the different management scenarios (STC = static closure; RTC = dynamic, real-time closure) for 2019 (left) and 2020 (right). The investigation used three indicators: the cumulative closure area and closure time, calculated by calculating the area under the graph by aggregating of the closure areas across 1 day (grey bars); the percentage of fishing mortality reduction (black triangles); and the fishing mortality reduction per cumulative closure area and closure time (red markers).

### 3.3. Economic effects

The STC performed at a lower level than all the RTC scenarios in terms of income achieved after opening the fishery following a closure (see Fig. 8). Analysis of the trend in the difference in income with and without closure, showed an observed gain in revenue if the area had been closed.

**Fig. 8.**
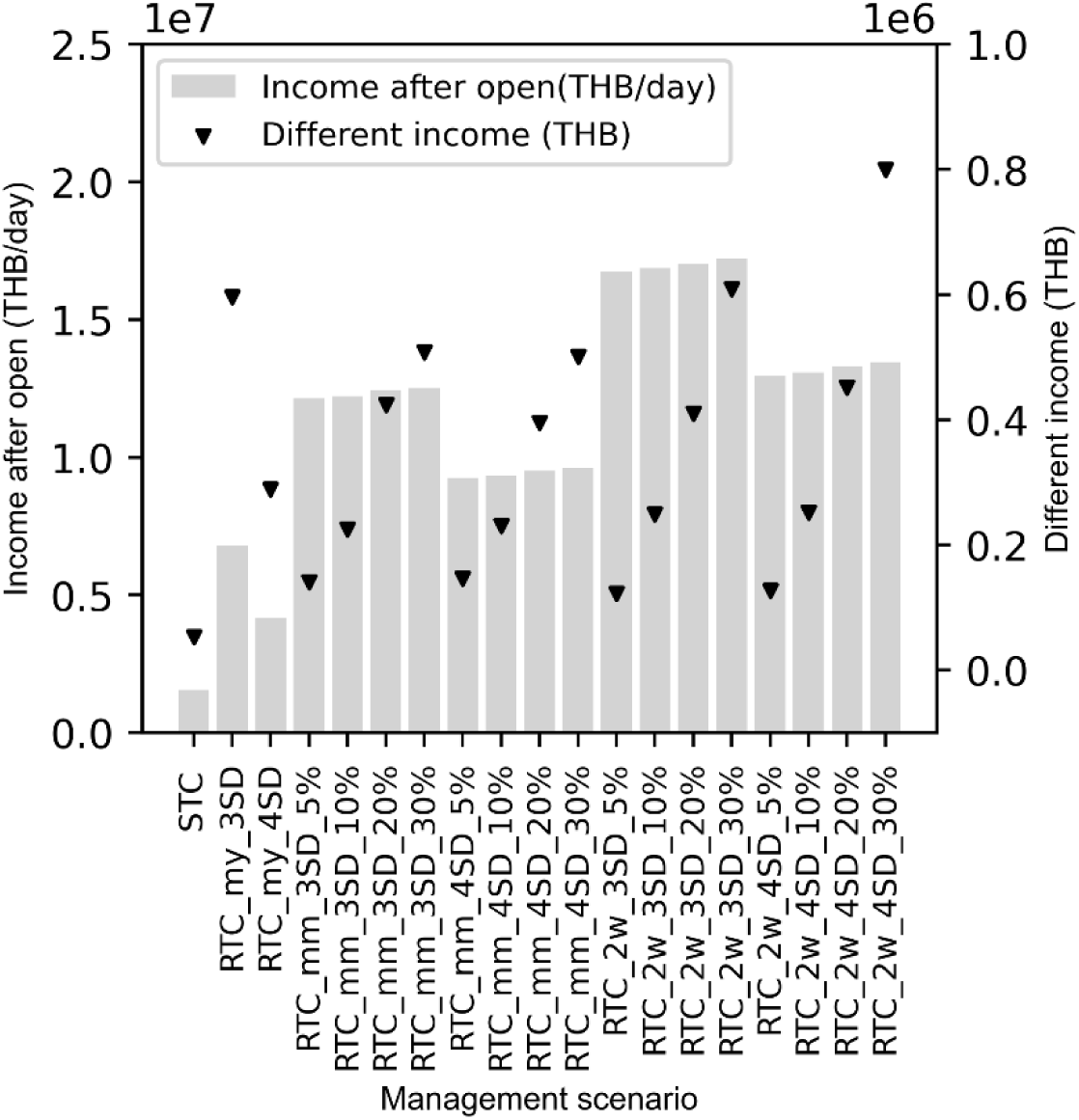
Performance indicators relating to economic effects in various management scenarios management scenarios (STC = static closure; RTC = dynamic, real-time closure), for 2019 (left) and 2020 (right). The two measures of performances were fishers’ income post-closure (Thai Baht [THB]/day) (grey bars), and the difference in income (THB) before and after opening (black triangles).

The implementation of closures resulted in higher income increases under an RTC when compared with a STC. When RTC scenarios were set by increasing the mortality reduction target percentage, the difference in income became larger, indicating a discernible trend (see Fig. 8). A comparative analysis of income after opening, per cumulative closure area and closure time, revealed that the income under the STC scenario was consistently lower than that under the RTC scenarios (Fig. 9).

**Fig. 9.**
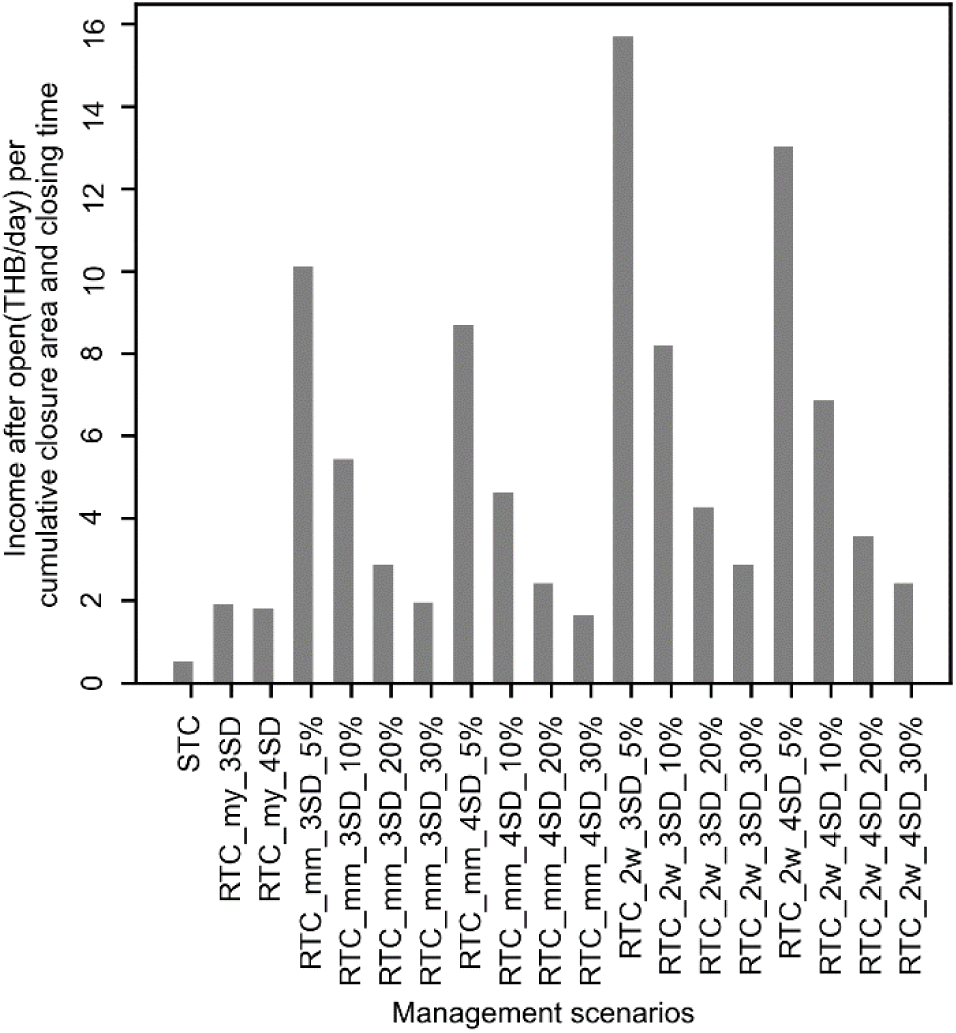
Comparison of fishers’ income after opening (THB = Thai Baht) shown against the cumulative closure areas and closure times in 2019 and 2020, under all management scenarios (STC = static closure; RTC = dynamic, real-time closure).

### 3.4. The potential of CPUE for defining the closure area

Our investigation of two CPUE thresholds (3SD and 4SD of the mean CPUE) to define the hotspot and closure unit under the RTC scenarios is illustrated in Fig. 10. No discernible disparity was observed in the percentage of mortality reduction between the two threshold levels (Fig. 10a). The mortality reduction per cumulative closure area and closure time was greater at the 4SD level than at the 3SD level (Fig. 10b). After opening the fishery closure, the revenue disparity exhibited a minor difference—the 3SD threshold went above the 4SD level in the triggered RTC (Fig. 10c). The analysis revealed that the income obtained after opening, per cumulative closure area and closure time, was greater at the 3SD than the 4SD level, especially for RTC scenarios with a mortality reduction target of 5% (Fig. 10d).

**Fig. 10.**
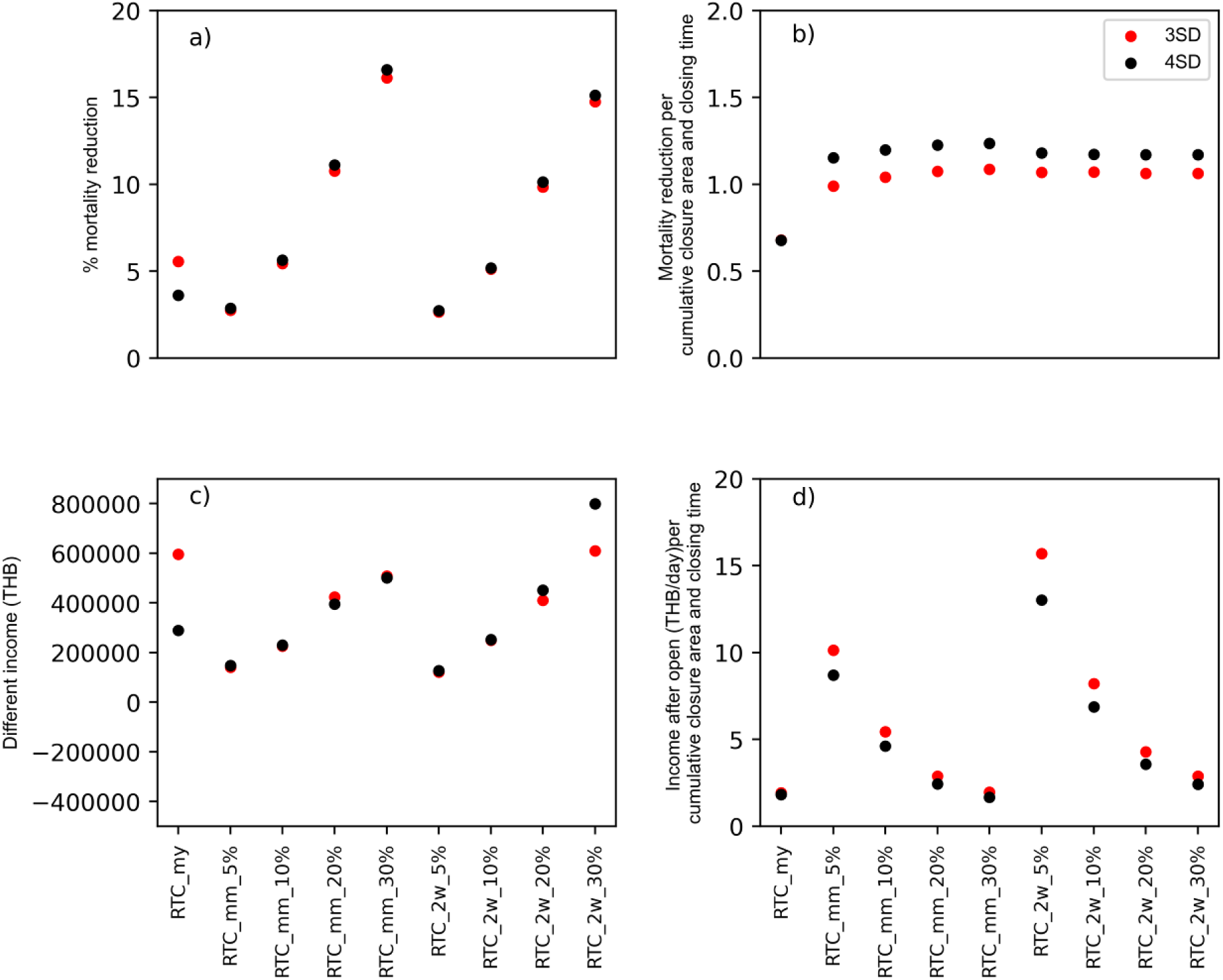
Comparison of closure performance at different CPUE thresholds, including 3 standard deviations (3SD, red dots) and 4 standard deviations (4SD, black dots) above the annual mean CPUE, used to determine hotspots for establishing units for real-time closures (RTCs) in the Gulf of Thailand in 2019. Closure performance was compared in terms of: a) percentage fishing mortality reduction; b) fishing mortality reduction per cumulative closure area and closure time; c) differences in fishers’ income (Thai Baht [THB]) before closure and post-closure; and d) fishers’ income (THB/day) after opening, per cumulative closure area and closure time.

### 3.5. Range of data collected for identifying CPUE and hotspots

Fig. 11 shows the variations in CPUE between the values obtained to create a hotspot for an RTC, and the values obtained during a closure, in relation to a data-collection interval of 1 year, 1 month, or 2 weeks. The results indicated that the time-range of data collection for detecting hotspots did not influence the differences in CPUE.

**Fig. 11.**
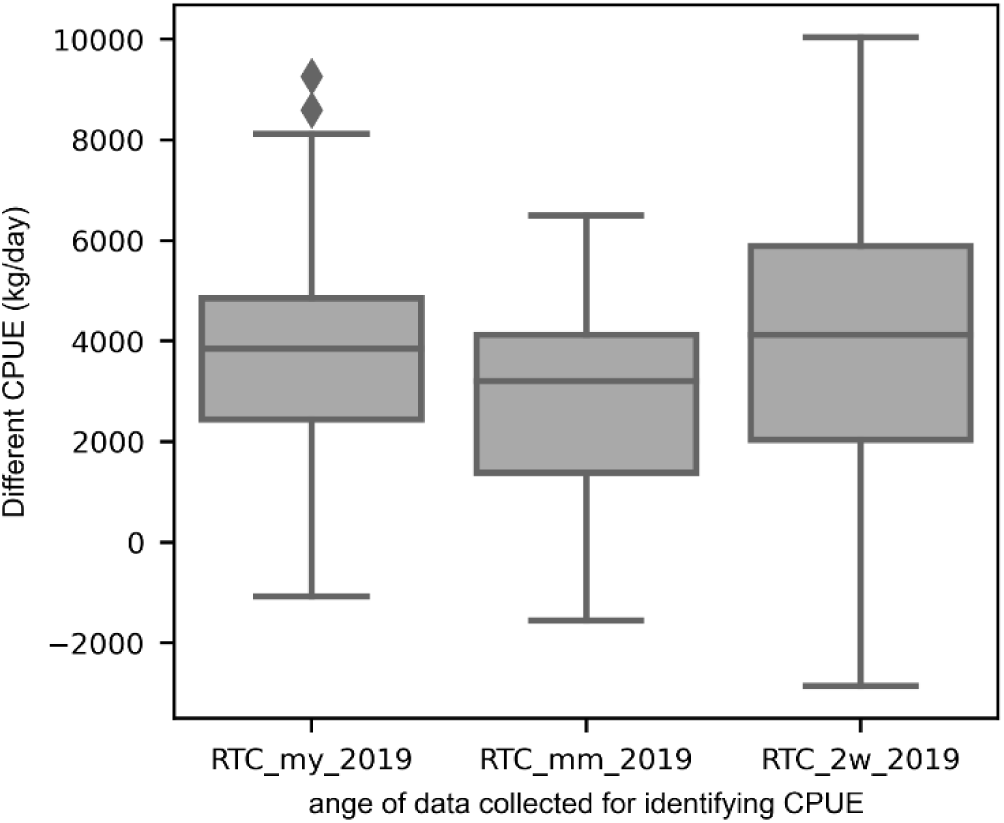
Boxplots of differences in catch per unit effort (CPUE) between predefined and post-determined hotspot closure units in the context of the real-time closure (RTC) scenarios in 2019. Data collection intervals: my = 1 year; mm = 1 month; 2w = 2 weeks.

## 4. Discussion

### 4.1. The STC versus RTC strategy

#### 4.1.1. Closure area and closure duration

Our analysis of the temporal relationship between the closure area and time (Fig. 5) confirmed that the STC strategy entails closure of the fishery at specific times during the calendar year, largely owing to mandated fisheries law, which was similar in the two years investigated here. Regarding the number of closed areas, the STC areas remain constant in every year.

The STC areas in the GOT are implemented in compliance with legislation that protects juvenile aquatic organisms during the spawning period, encompassing a broader range of species beyond short mackerel. Thailand’s five fishery closure areas are all near the coastline in the GOT and the ADS (Fig. 1). First, a monitoring system was employed to assess the suitability of the locations and durations of the STC. This was achieved by gathering data from commercial fishing vessels, research vessels, and biological information on the resources. Specifically, this involved evaluating the maturity status of bloodstock and the proportion of economically valuable fish larvae (Marine Fisheries Research and Development Division 2020). Consequently, the design of the STC has been driven by previous studies on migration patterns, emphasising spawning grounds. Afterwards, year-on-year measures were estimated to determine the appropriateness of implementing the area closures.

The RTCs showed discernible peaks in the total area closed to fishing during two specific time intervals, namely the 2nd and 4th quarters of the year; the area under the graph exhibited a narrow and peaked shape, which reflects closure in numerous management units. The closure times were also considerably lengthy during the 4th quarter. This trend became particularly evident when a 30% mortality reduction was required but fish density within the closed area was too low to meet the reduction target, requiring the closure to be extended into the following year. As a result, the graph appears relatively flat but prolonged, reflecting the extended time needed to achieve the 30% mortality reduction goal.

The conceptual framework of our RTC scheme uses the dynamic and degree of CPUE to define the management units that are to be temporarily closed to fishing. The number of units simultaneously closed was determined at levels 3SD and 4SD of overall CPUE. The duration of closure was based on the CPUE value (see equation in Section 2.3.2 and example in Fig. 4c). For instance, if the management unit being closed has a high CPUE, a short closure of a few days may effectively decrease fishing mortality in compliance with the planned target.

The implementation of an RTC focuses on monitoring the movement patterns of fish populations that are caught by commercial fishing fleets to establish temporary fishing closures in specific areas; the primary objective is to promptly identify areas of high fish abundance and implement closures to lower fishing mortality. The RTC areas exhibited a dynamic characteristic that fluctuated in response to fish migratory patterns. The temporal distribution of fishery closure areas exhibited a graph pattern characterised by smaller peaks and shorter closure durations, particularly from late in the 1st quarter to early in the 2nd quarter (Fig. 5) Implementing RTCs may have the potential to minimise the phenomenon of intense fishing effort near the boundary of a closed area, sometimes referred to as “fishing the line”. This phenomenon is frequently seen in the design of fixed seasonal closures, leading to overexploitation in waters surrounding the closed area (Farmer et al. 2020). The implementation of short-term closures like an RTC, and continual changes in the locations of the units and the overall size of the closure area, may ultimately help decrease the occurrence of fishing competition; however, the issue of fishing behaviour was not investigated in this study.

#### 4.1.2. Fishing mortality reduction

To achieve an equivalent reduction in the percentage fishing mortality, a STC requires greater closure area and closure time than an RTC. The effectiveness of reducing mortality under the STC approach was lower than that of the RTC approach in most scenarios, except for RTC scenarios where the mortality reduction target was 5% (Fig. 7). When comparing cumulative closure area and closure time, which represents the ratio of closure area to a unit of time, it was observed that the STC required closing a greater proportion of area and a longer closure time compared with an RTC. This difference was particularly notable for RTCs with a target mortality reduction of 5%.

The STC strategy primarily aims to implement fishery closures to protect small aquatic organisms and juvenile bloodstock. The RTC strategy likewise aims to reduce fishing-related mortality but without specifying size criteria for targeted fish. However, because RTCs achieve efficiency through their ability to establish a narrower geographic restriction and shorter temporal duration to minimise fishing mortality of target species relative to a STC, RTCs minimise lost fishing opportunities to capture other species.

Our study revealed that the effectiveness of closures to reduce fishing mortality depends on the distribution of the target species, and specifically the CPUE levels inside the closure units. The closure of a unit with a high CPUE is obviously effective in minimising fishing mortality. To explore this, our simulations included various scenarios wherein fishing mortality was set at distinct percentages (5%, 10%, 20%, or 30%). However, we found that the actual reduction in fishing mortality after closure was lower than expected (Fig. 7). Moreover, some units failed to meet the CPUE threshold to justify closure; in this case, we propose that when no unit achieves the CPUE threshold, it is preferable to proceed with closing whichever unit exhibits the highest CPUE value at that moment. Consequently, a closure area may be designated despite low CPUE, although this would compromise the efficiency of fishing mortality reduction.

These findings correspond to the research of Smith et al. (2021) which determined that the effectiveness of closure measures is dependent on both the spatial distribution of the species and the fishing activity. Even so, uncertainty about fish distribution may arise from various factors, including changes in the surrounding environment (Smith et al. 2021) or inappropriate methods used to predict fish distribution (Ducharme-Barth and Ahrens 2017).

Thus, it is imperative for future research initiatives to consider the application of RTC rules in situations where no particular unit achieves a threshold level of fish abundance (CPUE). In such a case, the need to implement a closure policy is questionable, if closure itself does not effectively mitigate fishing mortality of the target species and might also diminish opportunities to target other species.

#### 4.1.3. Economic impacts

The economic impacts of implementing a STC or RTC were considered in terms of two issues: comparison of a full-year’s income after opening the closed area (thus, only for units that experienced closure) (Fig. 8); and the difference in daily fishing income before implementing the closure and after opening it, for each respective scenario (Fig. 8).

Income after opening the STC was consistently lower when compared with that in all RTC scenarios, even though the cumulative closure area and closure time of the STC was greater than that of the RTCs, including the trigger area at the 4SD threshold level and with the target mortality reduction set at just 5% or 10% (Fig. 8). This difference in income might be attributable to low CPUE inside the closed areas as well as to the age composition of the fish population within that area. We used a population dynamic model (see section 2.4.) to examine the effect of age structure on fish population size and individual sizes when a fishing area is closed, because the income of fishers will be affected by a decrease in the number of fish.

The length frequency of short mackerel in the STC area, indicative of the age distribution of the fish population, suggested a high proportion of mature fish (mean ∼17.7 cm fork length [FL]). However, because closure times in the STC areas ranged from 60 to 90 days, we could presume a reduction in the overall number of fish because of natural mortality associated with older age. Furthermore, when a fish reaches maturity, its size does not necessarily increase considerably and its senescence begins, hence the quantity of fish will decrease over time owing to mortalities. This relationship is supported by growth parameters (*K*) and mortality estimates (Table 3), as well as trends in number-at-age (i.e. age structure) and yield (Fig. 12). This will accordingly have an impact on the income generated from a potential closure of the area.

**Fig. 12.**
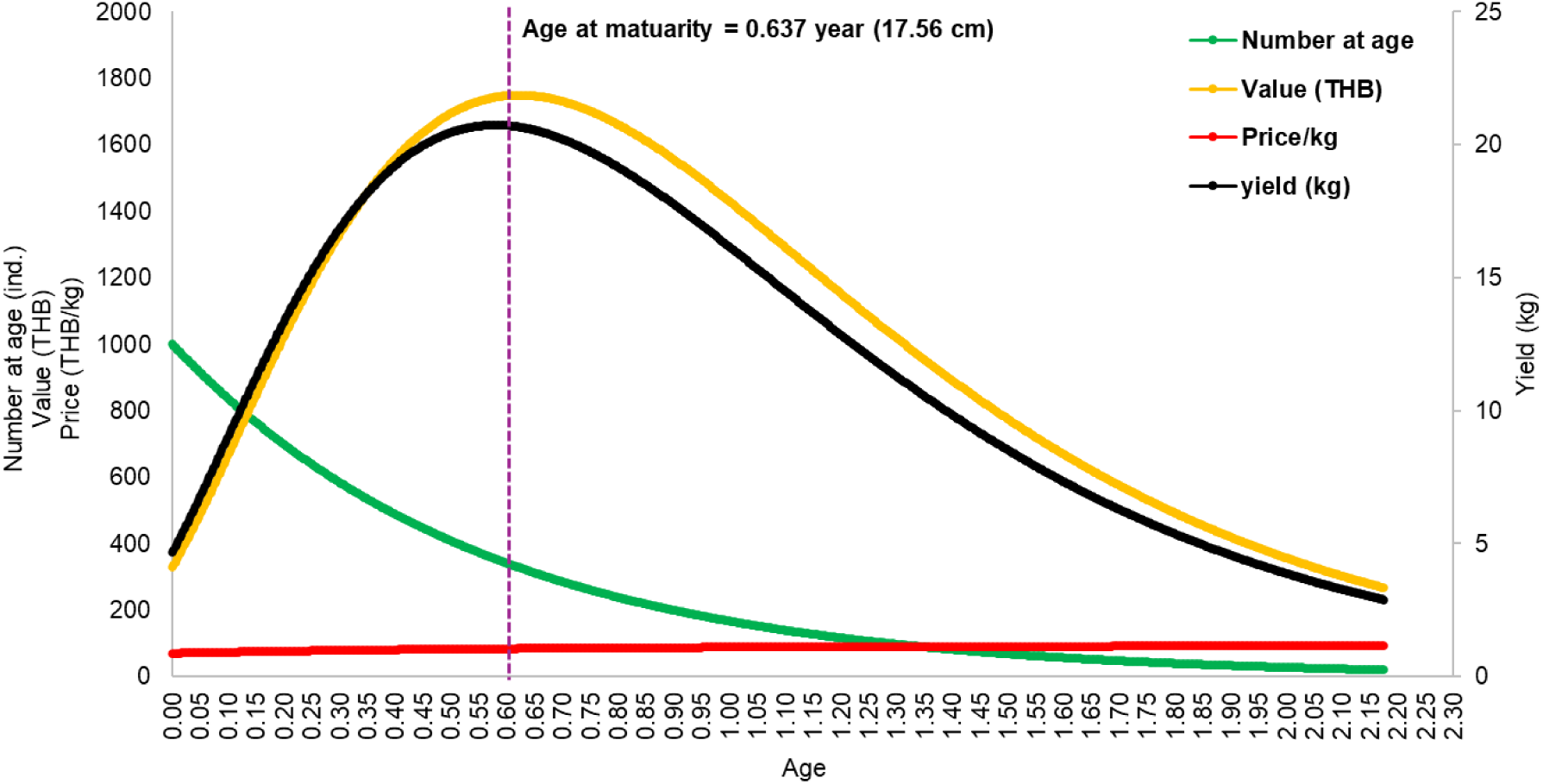
Relationship between the age (years) of short mackerel and other aspects of the catch in the Gulf of Thailand, including the number of fish caught, market prices, and value in Thai Baht (THB), using the same biological characteristics as in the management strategy evaluation. The number of fish (green line) was determined using the exponential decay model and the von Bertalanffy growth equation (Sparre and Venema 1998). Fish prices (red line) were approximated with a linear equation using fish pricing data from this research to make the graph smoother. Fish yield (black line) was estimated by multiplying the number of fish by the average weight of individuals (kg) at each age. The value in THB (yellow line) was estimated by multiplying the fish yields by the approximate market price at each age. The vertical, dotted purple line marks the age at maturity.

To calculate income, we determined the average prices of short mackerel in the fish market, and noted only a small difference in pricing between small fish (<17.5 cm FL, valued at 75.11 Thai Baht [THB]), medium fish (17.6–20.0 cm FL, 83.13 THB), and large fish (>20.0 cm FL, 96.29 THB). Consequently, closure of an area was unlikely to result in a meaningful rise in fishers’ incomes from landings of this species post-closure.

In contrast, the RTCs consistently exhibited more income post-closure compared with the STC across all scenarios. An examination of the length structure of the fish population within the RTC areas revealed a predominance of young individuals or immature fish. If these areas are closed to fishing, the size of short mackerel will increase over time based on the population dynamic model. Despite a general decrease in the number of fish in the population, it is notable that the yields consistently increased until reaching the maximum level when fish reached the length of maturity, as evident in Fig. 12. This phenomenon could be boosted by a closure of numerous units as part of the RTC management strategy. Even so, some units exhibited a juvenile age structure, as previously described. Additionally, the RTCs lasted for a relatively short time in each cycle. Consequently, the fish may not achieve maturity during the closure time as they enter a period when individual growth tapers off and the population decreases. This situation may result in greater overall income for fishers operating in an RTC compared with that for those operating in a STC after opening.

According to the population dynamics model for short mackerel applied in this study (see Fig. 12), fishery closures in areas with high juvenile abundance are expected to significantly improve overall production. We found that the size of fish (as weight) was increasing though the decline in the population had not yet reached a threshold that would negatively impact production, as depicted in Fig. 12. In this study, closure of an area and the outcome on characteristics of the fish age structure were surprising. Typically, an area closure might result in a loss of revenue, but in the case of the purse-seine short mackerel fishery in the Gulf of Thailand we determined that fishers’ incomes likely increased after a closure area was opened. This result suggests that if an RTC is designed in an appropriate location and time, it can benefit fishers in terms of their income; this fact could be leveraged to negotiate RTC agreements with fishers.

### 4.2. The CPUE threshold for defining the closure units

Two thresholds of CPUE were examined to define the closure region, 3SD and 4SD. We concluded that the appropriate CPUE threshold for determining a hotspot and closure unit is at the more restrictive 4SD because it entails less area and time, with minor variations in performance of the closures, particularly mortality reduction. Our results indicated that differing CPUE threshold levels did not influence the level of fishing mortality reduction nor the difference in fisher income before and after opening (Fig. 10). However, when considering the percentage of mortality reduction per cumulative closure area and closure time, the 4SD level exhibited greater decreases in fishing mortality than the 3SD level (Fig. 10b). This outcome was not unsurprising, given that the 4SD level commanded fewer closure units and less closure time.

The simulations showed that revenue after opening, relative to the cumulative closure area and time, was slightly higher at the 3SD than the 4SD level. This difference was particularly noticeable when fishing mortality reduction was set at 5%, whereas it varied only slightly at the other set percentages. This is attributable to a slight differential in the cumulative closure area and closure time between the two thresholds, even if the revenue generated after reopening a closed unit showed a greater contrast (Fig. 10d).

### 4.3. Appropriateness of data preparation for RTC implementation

In the context of this study, gathering data 1 month in advance was deemed appropriate as a sufficient interval to determine CPUE and detect hotspots to determine closure units, thereby offering the advantage of reducing the workload required for data collection. The difference between the CPUE value before defining the given closure unit and the CPUE value during a closure serves as an indication of the uncertainty associated with fish migration, reflecting the efficacy of RTC measures. The simulations conducted in this study included the immediacy of data collection appropriate for identifying CPUE hotspots and closure units. Specifically, three data-collection intervals were evaluated: 1 year, 1 month, and 2 weeks in advance (see section 2.3). No statistically significant variations in CPUE were observed across several scenarios involving different data-collection intervals (*p ≥* 0.05) (Fig. 11); boxplots summarising differences in CPUE between predefined and post-determined hotspot closure units show an observed CPUE discrepancy of approximately 3,594.03 across all scenarios (Fig. 11).

The 2-week data-collection interval in this study matches the data-gathering period used for an RTC strategy that aimed to mitigate cod fishing mortality under the Scottish Conservation Credits (SCC) scheme (Holmes et al. 2011), whereby VMS and fishing logbook data were collected over 14 days to identify areas for possible closure. However, the prior research did not explain the variations in CPUE observed after the closure. The ultimate objective of the SCC system is to attain an annual catch that effectively meets the mortality reduction targets given in a long-term management plan for cod. It is crucial to ensure that an RTC unit is designed so that CPUE remains consistently high throughout the first stages leading up to the closure, thereby mitigating the fishing mortality rate.

### 4.4. RTC implementation guidelines for use with existing surveillance systems

#### 4.4.1. Objectives and implementation

Dynamic management like RTC implementation is beneficial for conserving populations of migratory fish, which are characterised by a high level of unpredictability (Pons et al. 2022). The usefulness of closure identification is contingent on the precision of the approach used to ascertain CPUE via use of VMS data and landing reports, and information on other related variables. Inaccuracies might lead to identifying areas for closure in regions with low fish abundance, whereas locations with the greatest CPUE may be selectively dismissed (Holmes et al. 2011).

Although there is evidence that short mackerel are highly migratory in the GOT, some stocks in the eastern or western parts of the gulf are not migratory—as spatiotemporal distribution data showed no discernible movement trend over 3 years (Marine Fisheries Laboratory 1965). A species’ distribution will influence the performance of an area–time closure (Smith et al. 2021). If a closed area is designed to correlate with high fish abundance by area and time, then the closure might be highly successful.

Fisheries with robust data and management systems are appropriate for implementing RTCs, which rely on rapid information gathering and analysis. Hence, it is important to consider the accessibility of the data sources required to assemble comprehensive information on an area’s fishing activity and fish spatiotemporal distributions. The accuracy and precision of the information are equally crucial and will ultimately influence the performance of an RTC. The validation of fishing operation position data is an additional concern that warrants careful examination. Furthermore, the reliability of this information can be enhanced by incorporating supplementary camera technologies; for instance, using time-lapse technology on the deck of boats enables the observation of fishing activities. Confirmation of fishing activities can serve to validate the quality of the training data and enhances reliability of the information.

Electronic logbooks are another viable information source, and these can be integrated with the VMS position data by adding supplementary features to enhance the functioning of the VMS. The skipper has the ability to activate a button that serves to signal the start and end of the fishing activity. The geographical coordinates can be supplied to the report and utilised as an electronic logbook. Using electronic fishing operation location data can significantly reduce the data collection and processing time for analysis, thereby facilitating a more accurate determination of fishery closure areas.

To use RTCs to minimise fishing mortality of a target species without defining the age range, the surveillance data could potentially be sufficient to support the decision to implement a closure. Furthermore, the RTC scheme we describe may be conveniently modified to accommodate different species that exhibit varying levels of uncertainty in their migratory patterns. In the present investigation, gathering data 1 month in advance appeared sufficient to implement an RTC to benefit short mackerel. This seems a reasonable time-frame for staff to gather information and announce an area closure because our investigations found no variation in CPUE at this data-gathering interval under the different scenarios. Additionally, 2 weeks may be insufficient for fishers to organise a trip. However, if extending this analysis to other fish species characterised by higher levels of uncertainty, biweekly or weekly data-collection intervals could be considered to identify the locations of hotspots. Additionally, the closure units could vary in size, either bigger or smaller, compared with the ones we proposed.

If the primary goal of an RTC extends beyond a reduction in fishing mortality, specifically in terms of mitigating the mortality of bloodstock or immature fish, its implementation may have different economic effects. For instance, implementing an RTC to protect juvenile fish might potentially lead to an increase in revenue after the closed area is reopened.

As mentioned, the application of RTCs is a complex process that necessitates the rapid acquisition of substantial information. This includes data preparation, such as collecting relevant information to identify areas with high fish abundance, which then informs the establishment of closure units. Furthermore, it requires law enforcement agencies to possess efficient monitoring and surveillance tools, such as implementing VMS by fisheries monitoring centres, to accomplish the objectives of the RTC initiative effectively. Hence, it is recommended that RTCs be implemented for fish populations that are assessed as being in a critical state. RTCs represent a highly successful management intervention owing to their notable effectiveness in reducing fishing mortality and capacity to accomplish management objectives within a short time-frame.

#### 4.4.2. Enforcement

Enforcement is a considerable challenge to fishery closures. Three potential approaches might be considered: voluntary engagement, co-management, and governance enforcement. RTCs have been previously implemented through several approaches, primarily determined by the characteristics of the fisheries and the existing legislative framework. The RTCs to manage short mackerel in Thailand have been successful partly because of the recent enactment of fishing legislation that facilitates the implementation of area closures; hence, a top-down strategy with government enforcement has favoured this situation. Relevant fisheries laws in Thailand’s Royal Ordinance on Fisheries B.E. 2558 (2015) (Department of Fisheries Thailand 2015) concern time–area closures, including Section 70 which states: “*No person shall engage in a fishing operation during a season of aquatic animals’ ovulation and egg-spawning, larvae rearing or during any other period designated for the protection of aquatic animals as prescribed by the Minister*”.

Furthermore, Thailand has established a VMS for fishing operations inside restricted zones; thus, this surveillance task has been incorporated into the regular operations of the Fisheries Monitoring Centre (FMC). The current VMS monitoring web system has an organisational layout for restricted areas, with FMC officials now providing 24-h monitoring to ensure continuous surveillance.

Nevertheless, RTCs uniquely require periodic updates on the enclosed areas. Hence, it may be necessary to develop an application that enables the modification of restricted configurations within the VMS to display via the web application, facilitating the monitoring capabilities for both fishers and FMC officials.

#### 4.4.3. Challenges of RTCs

The unpredictability of fish migration patterns is a consistent issue in the design of time–area closures. Differences in CPUE between the time data are gathered and a closure is implemented are likely to occur regardless of the data-gathering period—whether it takes 1 year, 1 month, or 2 weeks. In this study, after the initial data were gathered to determine the closure units, it was discovered that the CPUE in some units fell to zero during the closure. Hence, establishing a hotspot with a predetermined duration requires considerable thought because it directly impacts the efficacy of the closure. Ongoing monitoring of fish movements is important because the periodic nature of these patterns may differ from year to year.

The implementation of RTCs involves significant difficulties and challenges. A first concern is their acceptance by fishers, while a second concern is the complexity of their implementation and enforcement. Although it is advisable to use top-down strategies for the implementation of an RTC, agreement must exist among stakeholders. In negotiations with fishers, it is helpful to highlight the financial effects associated with the depletion of fish populations in areas of abundance, such as juvenile fish.

Another significant concern is the provision of prompt notification to fishers about the accessibility of closure units. In some nations, updates are disseminated via announcement websites and radio broadcasting (Little et al. 2015). In the context of Thailand’s fishing industry, radio communication is especially helpful for delivering information on RTC units. Closure areas are typically square, which makes it easy to identify a unit’s borders, although a substantial number of precise coordinates must be provided. Thus, using the VMS online application to update RTC units is prudent. Typically, fishing vessels equipped with a VMS can monitor their own geographical coordinates. Exploiting this tool would afford fishers an alternative channel for obtaining information about closure of units.

The simulations used in our research did not investigate the complexities associated with multi-species dynamics. However, it was evident that the RTCs effectively mitigated the fishing mortality of short mackerel, which we confirm is present a single species in the study areas. The implementation of RTCs to protect a specific fish species from fishing mortality raises the question of whether considering multiple species in closure design might end up diminishing economic returns and interrupting fishing opportunities for other captured species. Further investigation is required to demonstrate the effectiveness of RTCs to conserve a range of fish species of different age distributions, and in connection with different habitats. Future research should seek to clarify the potential and limitations of implementing multi-species spatial management strategies.

## 5. Conclusions

This research clarifies the efficacy of using dynamic RTCs instead of static fishery closures to minimise fishing mortality, while also exploring the utility of surveillance data on migratory fish to design effective RTCs. The RTC management approach offers more flexibility and requires less closure area than implementing a STC in terms of the efficiency of the same units to achieve management objectives via measures such as minimising mortality or increasing fishers’ income after opening the unit. Furthermore, it is also feasible to establish a targeted reduction in fishing mortality based on the CPUE threshold. To maximise the economic benefit from implementing an RTC, it is essential to thoughtfully arrange closures of areas with abundant juvenile fish.

We recommend employing the highest CPUE threshold level to define a closure unit and gathering data on CPUE monthly to alleviate the administrative burden. Furthermore, no management unit should be shut down until all units achieve the CPUE threshold level. Further study should be undertaken to explore the integration of age structure data, particularly data obtained the previous year, as was combined with near real-time surveillance data in this study.

In the future, integrating data beyond surveillance information, such as fish size data, could further inform the design of area closures. This approach may support a broader range of management objectives, including closures aimed specifically at reducing mortality of juvenile fish.

## Acknowledgements

We express our sincere thanks to Thailand’s Department of Fisheries and to all the administrative staff who provided fishing logbooks, VMS data, fishing trip records, and landing information, which was essential for conducting the analysis. Cynthia Kulongowski with Edanz (https://jp.edanz.com/ac) edited the language of a draft of this manuscript.

## Declarations

### Data availability

All data sources used in this study are cited within the manuscript. Surveillance datasets, including the Vessel Monitoring System (VMS), trip records, logbooks, and landing reports, were obtained with formal authorisation from the Thailand Department of Fisheries (DOF). These datasets contain confidential information, including vessel identification numbers and personal data, and are protected under the Royal Ordinance on Fisheries B.E. 2558 (2015) and related regulations. For the purpose of analysis, the DOF supplied anonymised proxy vessel identification codes to enable vessel-level classification without disclosure of personally identifiable or sensitive information. Owing to statutory confidentiality and data-protection obligations, the original raw data are not publicly available and may only be accessed subject to prior approval and applicable legal permissions from the Thailand Department of Fisheries.

### Author contributions

**CM:** Conceptualised the study, developed methodology, coded and implemented the model, analysed data, visualised results, and drafted the manuscript. **PN:** Refined methodology, curated and validated data, interpreted results, and reviewed the manuscript. **PS:** Assisted with data collection, preprocessing, analyses, and technical review. **WS** and **MK:** Provided expertise in Thai fisheries policy and management and contributed to discussion and policy implications. **TY:** Supported software development, model validation, and verification of analytical outputs. **MTF:** Supervised the project, advised on study design and analysis, and reviewed and edited the manuscript.

